# Manipulation of the Extracellular Vesicle Biogenesis Pathway in Paternal Somatic Cells Contributes to the Phenotypic Fate of the Next Generation

**DOI:** 10.1101/2025.04.07.647377

**Authors:** Emma Louise Salt, Enzo Scifo, Kristina Schaaf, Maryam Keshavarz, Ting Liu, Xieze Wei, Daniele Bano, Dan Ehninger

## Abstract

Human and animal studies have shown that risk factors for certain diseases are linked to our father’s physiological and psychological condition. So far, mainly *in vitro* evidence indicates extracellular vesicles (EVs) as one of the potential mediators of these paternal effects. In a full *in vivo* model system, we demonstrate that targeting the EV biogenesis pathway within somatic cells of the male *Drosophila melanogaster* reproductive tract induces sex-specific phenotypic changes in the progeny. These findings highlight the critical role of the EV pathway in paternal somatic cells in shaping the phenotypic destiny of the next generation and underscore the necessity for further investigations into the mechanisms of somatic to germline information transfer.

## Main Text

The ‘Weismann barrier’ hypothesises that heritable information cannot be transmitted from somatic cells to germ cells, suggesting the phenotypic fate of our offspring is determined purely by the luck of the Mendelian genetic draw [1]. With the rising awareness of the impact experiences and life style choices have on our health, it remains to be determined whether these factors can also affect our descendants and if so, what the molecular mechanisms are. Evidence from human and animal studies indicates that changes to the parental physiological or psychological state can lead to increased risk for pathologies such as metabolic disorders, cardiovascular disorders, cancers and psychiatric disorders in the next generations [2]. Although society’s focus is mostly on the impact of the mother, these so-called intergenerational effects can stem also from the father and are observed across various species, from *Homo sapiens* to *Drosophila melanogaster* [3-5]. A range of mechanisms including changes to the sperm epigenome – for example, DNA methylation, histone modifications and sperm RNA content changes – have been identified to contribute to these paternal effects [6]. How paternal experiences, conditions or exposures cross the ‘Weismann barrier’ remains unclear.

Extracellular vesicles (EVs) secreted by somatic cells have been proposed to transmit information to the germ cells, leading to changes in the phenotypic outcome of the next generation [7, 8]. These small membrane-bound particles are released from nearly every cell and function as intercellular communicators through their transport of cargos, such as DNA, proteins and various RNA species, to other cells [9]. In the male reproductive tract, EVs have been shown to transmit proteins or RNA to the sperm and are essential for sperm maturation [10-12]. So far, available evidence to support a role of somatic cell-derived EVs on the phenotypic fate of the next generation is limited to studies using *ex vivo* manipulations of sperm [8]. Here, we introduce the use of a full *in vivo* model as an experimental approach to analyse this relationship between somatic cell-derived EVs and the offspring phenotypic outcome.

Within the male reproductive tract of *Drosophila melanogaster*, the accessory gland (AG) generates many components of the seminal fluid, similar to the mammalian prostate [13]. At the tip of the AG, secondary cells (SC) have been shown to secrete EVs into the AG lumen [14] (Figure 1A). Upon mating, the seminal fluid containing seminal fluid proteins, sperm and EVs are transported into the female reproductive tract, where the EVs interact with germ cells [14].

**Figure 1:**
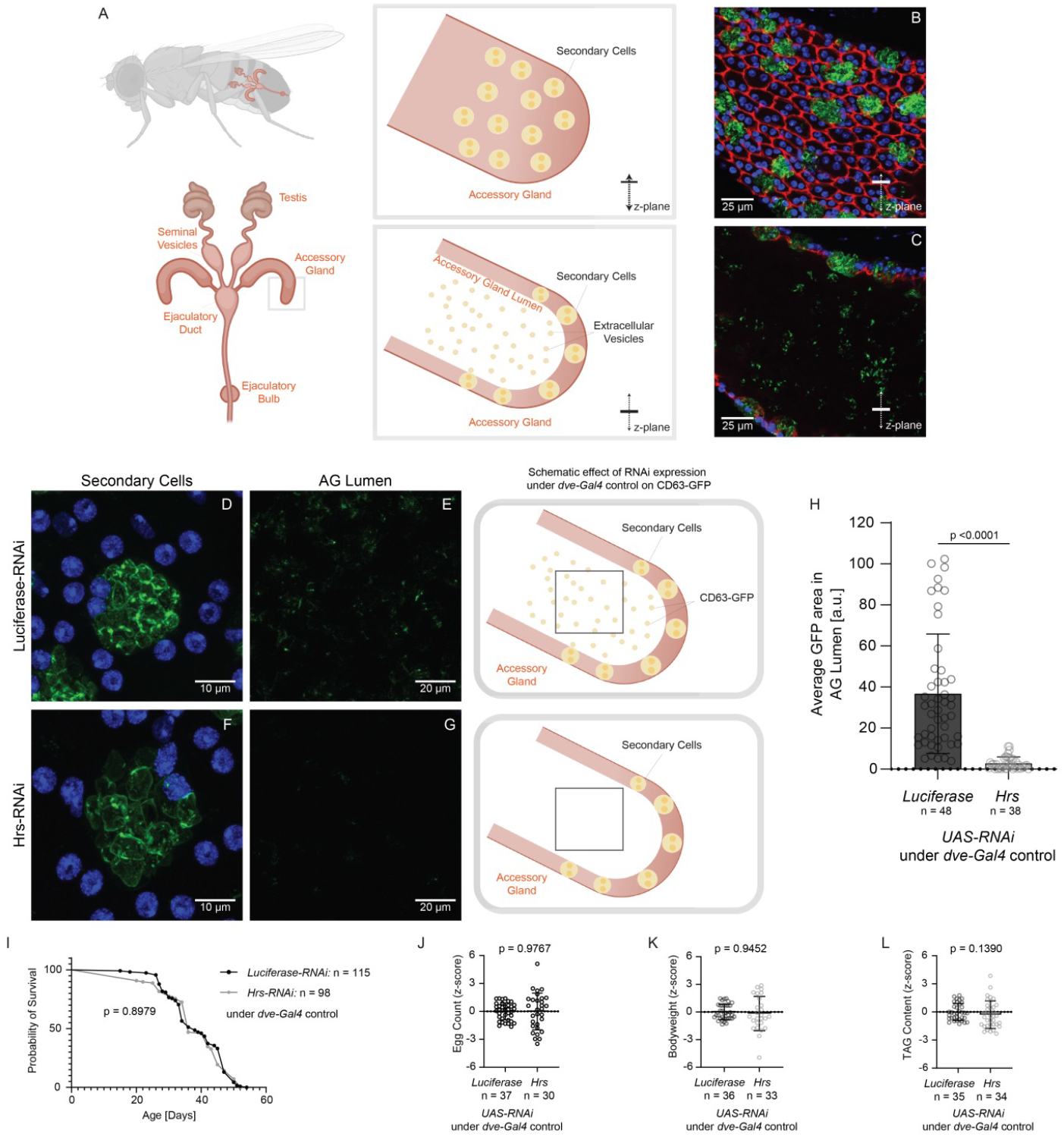
*Drosophila melanogaster* lines expressing RNAis under *dve-Gal4* control as an *in vivo* EV loss of function model. Secondary cells (SCs) within the tip of the accessory gland of the male *Drosophila melanogaster* reproductive tract secrete extracellular vesicles (EVs) into the accessory gland lumen, as previously described in [14] and depicted in the schematic diagram created using Biorender.com (A). FasIII (red) staining visualises cell membranes (B-C) and scale bars are shown in each image for size reference (B-G). *UAS-CD63-GFP* (green) is expressed in SCs under *dve-Gal4* control and localizes in their intraluminal structures (B). Outside of SCs, CD63-GFP puncta are visible in the accessory gland lumen (C). The expression of *Luciferase-RNAi* (D and E) and *Hrs-RNAi* (F and G) under *dve-Gal4* control leads to visible changes in the intraluminal structures of SCs and the number of CD63-GFP puncta within the accessory gland lumen. Quantification of GFP positive pixels in defined sections of the accessory gland lumen in flies expressing *Luciferase-* and *Hrs-RNAi* under *dve-Gal4* control is displayed in bar graphs with the mean ± SD indicated and individual data points superimposed (H). The lifespan of flies expressing *Luciferase-* and *Hrs-RNAi* under *dve-Gal4* control is plotted with the Kaplan-Meier method and statistically analysed using Log-rank (Mantel-Cox) test with p-value displayed in graph (I). Z-standardised number of eggs laid by female partners (J), bodyweight (K) and TAG content (L) of male flies expressing *Luciferase-* and *Hrs-RNAi* under *dve-Gal4* control is shown in bar graphs with mean ± SD indicated and individual data points superimposed. Unpaired t-test with Welch’s correction or Mann Whitney test was carried out for statistical analysis, with p-value displayed in graph (H, J-L). Sample sizes are displayed in the respective graphs and refer to a pool of 5-10 flies in panel K and 5 flies in panel L. SD = standard deviation.

As a common component of EVs, labelling of CD63 is a well-established EV marker that has been employed in several publications [14,15]. In this study, we express human CD63-GFP under *dve-Gal4* control using the UAS-Gal4 system. In the reproductive tract of adult male *Drosophila melanogaster, dve* is expressed highly in SCs (Figure 1B,C). Within SCs, CD63-GFP localises in intraluminal structures (Figure 1B), while CD63-GFP puncta are visible externally of the SCs within the AG lumen (Figure 1C). Under the same *dve-Gal4* driver, we generated *Drosophila melanogaster* lines that express either *Hrs-RNAi* (targeting a known component of one of the EV biogenesis pathways) or *Luciferase-RNAi* (targeting a firefly gene, as the control) (Figure 1D-G). CD63-GFP localizes in intraluminal structures within the SCs of both genotypes (Figure 1D,F), while CD63-GFP puncta are barely visible in the AG lumen of male flies expressing *Hrs-RNAi* under *dve-Gal4* control (Figure 1E,G). The quantification of the GFP area within defined sections of the AG lumen confirms the observation that the expression of *Hrs-RNAi* under *dve-Gal4* control significantly decreases the secretion of CD63-GFP into the AG lumen (Mann Whitney test: p < 0.0001) (Figure 1H). Notably, no significant differences are observed in the lifespan (Log-rank test: p = 0.8979) or the number of eggs laid by female partners (Unpaired t-test with Welch’s correction: p = 0.9767) between the two genotypes (Figure 1I,J), indicating that the expression of *Hrs-RNAi* under *dve-Gal4* control does not lead to long-term health or fecundity impacts in the male flies. To check for changes in body composition, we assessed bodyweight (Mann Whitney test: p = 0.9452) and triglyceride (TAG) content (Mann Whitney test: p = 0.1390), both of which show no significant changes caused by the expression of *Hrs-RNAi* under *dve-Gal4* control (Figure 1K,L).

To investigate the impact of the genetic manipulation of the EV biogenesis pathway in paternal somatic cells on the phenotypic outcome of the next generation, male flies expressing *Hrs-RNAi* and *Luciferase-RNAi* under *dve-Gal4* control (F0) were crossed with wild type females and their 5-day old F1 offspring were analysed (Figure 2A). We selected F1 flies to carry identical balancers inherited from their fathers, such that all offspring included in our analysis did not inherit any of the paternal *dve-Gal4* or *UAS-RNAi* components. Two-way ANOVA identifies a significant paternal genotype effect on the bodyweight of young adult F1 flies (F (1, 81) = 11.37; p = 0.0011), with post hoc analysis showing a significant decrease in the bodyweight of male offspring from fathers expressing *Hrs-RNAi* under *dve-Gal4* control compared to *Luciferase-RNAi* offspring (Tukey’s multiple comparisons test: p = 0.0162). Young adult F1 females demonstrate a non-significant bodyweight decrease (Tukey’s multiple comparisons test: p = 0.3190) (Figure 2B). A significant paternal genotype effect is also observed in the TAG content of F1 flies with two-way ANOVA (F (1, 105) = 13.69; p = 0.0003) (Figure 2C). Tukey’s multiple comparisons test indicates a significant increase in the TAG content of male F1 flies from F0 fathers expressing *Hrs-RNAi* under *dve-Gal4* control (p = 0.0015), but no significant changes in the female F1 (p = 0.4428) (Figure 2C).

**Figure 2:**
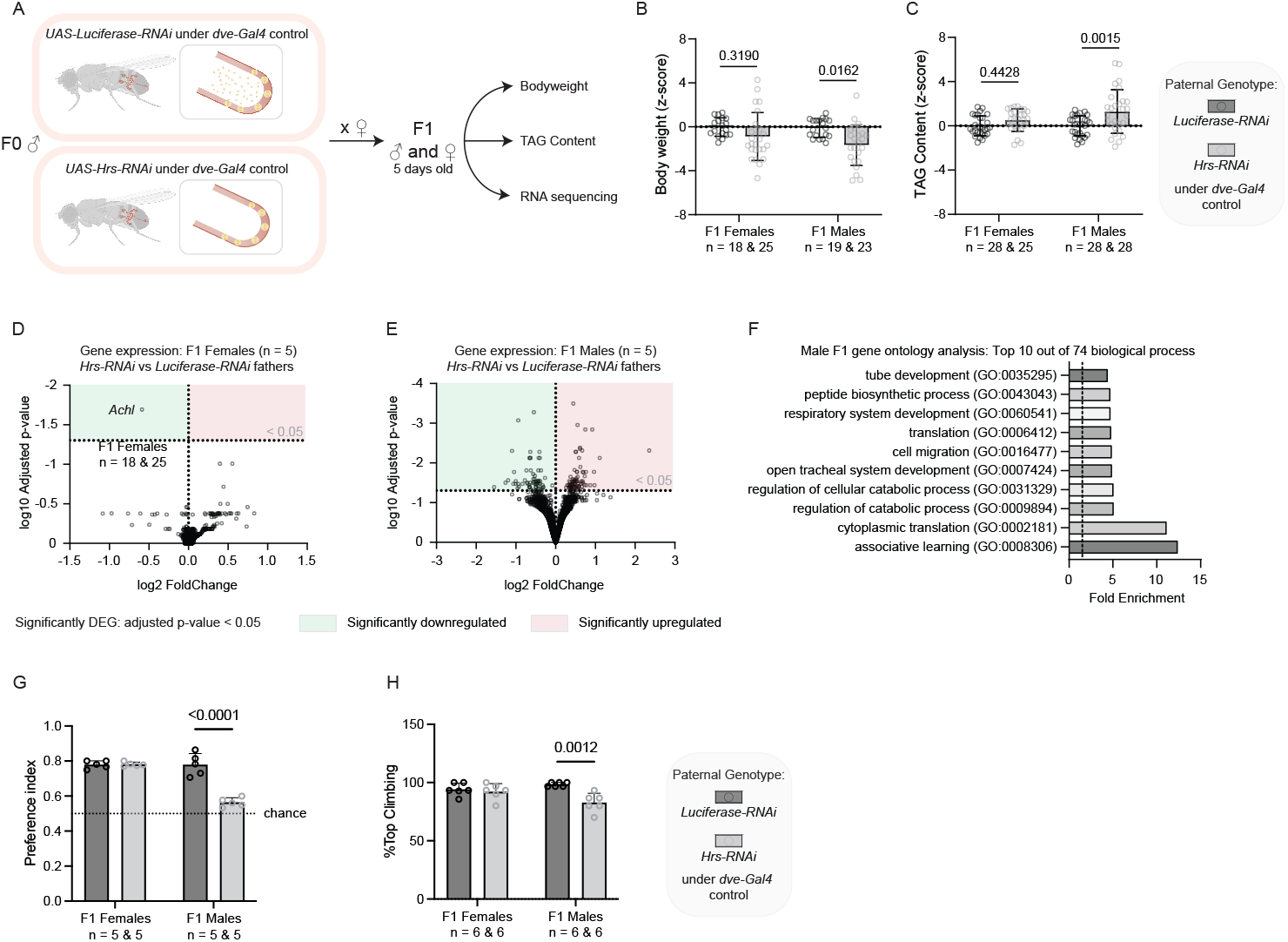
The phenotypic outcome of male and female F1 offspring from F0 male flies expressing *Luciferase-RNAi* and *Hrs-RNAi* under *dve-Gal4* control. Schematic diagram created with Biorender.com depicts F1 generation and analysis pipeline (A). Z-standardised F1 bodyweight (B) and TAG content (C) are displayed in bar graphs indicating mean ± SD (individual data points superimposed), with two-way ANOVA carried out for statistical analysis and p-value of Tukey’s multiple comparisons test shown within sex. Gene expression of female (D) and male (E) F1 offspring from *Hrs-RNAi* fathers vs *Luciferase-RNAi* fathers are displayed based on fold change and adjusted p-value on the logarithmic scale in volcano plots. The threshold for significantly differentially expressed genes (DEG) is the adjusted p-value < 0.05. Top 10 biological processes identified by gene ontology analysis (PANTHER, false discovery rate < 0.05) from significantly differentially expressed genes of F1 males are displayed based on fold enrichment (F). Performance on a T-maze-based associative learning task (G) as well as a climbing assay (H) are represented in bar graphs indicating mean ± SD (individual data points superimposed), with two-way ANOVA carried out for statistical analysis and p-value of Tukey’s multiple comparisons test shown within sex. Sample sizes are displayed in the respective graph and refer to a pool of 5-10 flies in panel B, 5 flies in panel C-E, 20-30 flies in panel G and 10 flies in panel H. SD = standard deviation.

To further study these sex-specific findings, the male and female offspring were analysed on the transcriptomic level. In young adult females from fathers expressing *Hrs-RNAi*, RNA sequencing reveals only 1 significant differentially expressed gene (DEG) in young adult females from fathers expressing *Hrs-RNAi*, the gene *Achl* (adjusted p-value < 0.05) (Figure 2D). In comparison, 145 significant DEG are identified in young adult male F1 offspring of *Hrs-RNAi* fathers compared to male offspring from F0 flies expressing *Luciferase-RNAi* under *dve-Gal4* control (Figure 2E). From these, gene ontology (GO) analysis identifies ‘associative learning’ (GO:0008306), ‘cytoplasmic translation’ (GO:0002181) and ‘regulation of catabolic process’ (GO:0009894) among the significantly enriched biological processes (adjusted p-value < 0.05) (Figure 2F).

Given these transcriptional alterations, we next examined associative learning and memory abilities of *Hrs-RNAi* F1 offspring. Towards this end, we used a T-maze to train flies to associate one of two specific odours (3-octanol, 4-methylcyclohexanol) with a sugar reward. Subsequent to training, we tested associative memory assessing odour preference (for details, see Methods). Our analyses demonstrate an overall substantially decreased preference index (PI) in *Hrs-RNAi* F1 offspring relative to controls (two-way ANOVA: F (1, 16) = 35.96; p < 0.0001) (Figure 2G). Posthoc analyses showed a significantly reduced PI in male mutant flies (Tukey’s multiple comparisons test: p < 0.0001), but not in females (Tukey’s multiple comparisons test: p = 0.9504), again in line with the notion of sex-specific influences of the paternal genotype on F1 offspring flies. Additional behavioural analyses, using a climbing assay, revealed further evidence for neurobehavioral changes in *Hrs-RNAi* F1 offspring: A paternal *Hrs-RNAi* genotype led to significantly reduced climbing abilities relative to controls (two-way ANOVA: F (1, 20) = 12.86; p = 0.0018) that was driven by changes in male (Tukey’s multiple comparisons test: p = 0.0012) but not female (Tukey’s multiple comparisons test: p = 0.9396) offspring flies (Figure 2H).

Overall, this study is one of the first to link the manipulation of the paternal somatic cell EV biogenesis pathway and the offspring’s phenotypic health in a complete *in vivo* model. We observe that manipulations that target the EV biogenesis pathway within somatic cells of the male reproductive tract using genetic tools can affect the phenotypic outcome of the next generation in a sex-specific manner. While sex-specific changes are not uncommon findings in paternal inter- and transgenerational studies [for example as seen in 16], we are the first to observe that genetic manipulations of the EV biogenesis pathway from somatic cells within the male reproductive tract can affect the phenotypic outcome of the next generation in a sex-specific manner. Ultimately, the detailed mechanisms leading to observed intergenerational effects, and whether EVs act alone or are a part of an elaborate signal cascade involving other of yet unknown co-factors are still to be determined and are beyond the scope of this study. Going forward, we encourage the use of this model and similar *in vivo* systems, in order to investigate EVs in soma to germline communication and their role in offspring phenotypic fate in *in vivo* settings.

## Acknowledgements

We thank Prof. Dr. Gaia Tavosanis and the members of her research group for the use of their laboratory space and support. We thank Prof. Tavosanis, Suzanne Eaton and Bloomington Drosophila Stock Center for flies and Developmental Studies Hybridoma Bank for antibodies. Furthermore, we would like to thank Prof. Joachim L. Schultze, Prof. Marc Beyer and Dr. Lorenzo Bonaguro from the PRECISE Team, DZNE e.V. Bonn for their support with the transcriptomic data. We are also grateful to Dr. Hans Fried and Dr. Ireen König from the Light Microscope Facility (LMF), DZNE e.V. Bonn for their support with the imaging acquisition.

## Author Contributions

DE conceived the research project and provided supervision; ELS, ES, KS, MK, TL and DE planned the research project; ELS, ES, KS, MK, XW and TL established assays and performed experiments; ELS, ES, KS, MK, TL and DE analysed data; ELS, ES, KS, MK, TL, DB and DE contributed interpretation and discussion; KS provided technical support; ELS and DE drafted the manuscript; DB and DE provided resources. All authors agreed on the final version of the paper.

## ‘Inclusion & Ethics’ Statement

The authors declare no competing financial interests.

## Methods and Materials

### Fly stocks and husbandry

The following fly stocks were acquired from Bloomington Drosophila Stock Center: *dve-Gal4* (w[1118]; P[w[+mGT]=GT1] dve[BG02382]/CyO - RRID:BDSC_12859), *UAS-Luciferase-RNAi* (y[1] v[1];; P[y[+t7.7] v[+t1.8]=UAS-LUC.VALIUM10]attP2 - RRID:BDSC_35788), *UAS-Hrs-RN*Ai (y[1] sc[*] v[1];; P[y[+t7.7] v[+t1.8]=TRiP.HMS00841]attP2 - RRID:BDSC_33900). UAS-CD63-GFP line (w[*]; P[w[+mC]=UAS-EGFP.CD63]2; Dr[1]/TM3, Sb[1]) was donated by Suzanne Eaton, Max Planck Institute of Molecular Cell Biology and Genetics. All stocks were crossed with double balancer (yw;IF/CyO;MKRS/TM6B) to generate lines with comparable balancers. Flies were kept at 25°C and 60 % humidity with a 12h/12h light/dark cycle on a standard food diet.

### Imaging of the AG within the male *Drosophila melanogaster* reproductive tract

The whole reproductive tract of male flies expressing *UAS-CD63-GFP* and *UAS-RNAis* under *dve-Gal4* control was dissected in cold 0.03 % PBST. Following fixation in 4 % PFA, if applicable, staining was carried out by permeabilising the tissue with 0.3 % Triton X-100 and blocking with BSA. The tissue was then incubated overnight at 4 °C with the primary antibody against Fascilin III (7G10 anti-Fasciclin III was deposited to the DSHB by Goodman, C.; 1:10), with another 2 hours incubation at room temperature with a secondary antibody carrying Alexa Fluor 594 (Life Technologies A11005, 1:400). The tissue was mounted on microscope slides using VectorShield mounting medium containing DAPI (Fisher Scientific #13273694). Images were acquired using a LSM700 confocal microscope with 40x and 63 x objectives, and in some cases 2x zoom. Representative images are shown as maximum intensity projections. Gain and laser intensity settings were optimised for the image acquisition.

### Quantification of CD63-GFP within the *Drosophila melanogaster* AG lumen

Confocal images of the AG lumen were acquired with a 63x objective and using standardised gain and laser intensity settings between samples for quantification. Imaging was carried out from the top to the bottom epithelial cell layer of the AG in 3 different sections in 1 µm slice intervals. Starting after the first 10 confocal slices, the total GFP area of 20 slices (starting after the first 10 slices to exclude the epithelial cell layer) was quantified in arbitrary units [a.u.] and averaged per section and per AG. The following number of samples were removed as outliers from each dataset using ROUT (Q = 1 %): Luciferase-RNAi n = 1, Hrs-RNAi n = 3.

Sample sizes of imaged AGs for the CD63-GFP puncta quantification were as follows: *Luciferase-RNAi* n = 48, *Hrs-RNAi* n = 38.

### Lifespan analysis

Fly handling and lifespan analysis were carried out as previously described by Linford, Bilgir [17]. In short, male F0 flies were flipped into fresh vials every 2-3 days with dead, escaped or stuck flies documented every 2 days. Data was plotted using Kaplan-Meier survival curves and analysed with a Log-rank (Mantel-Cox) Test.

The sample sizes for the lifespan analysis were as follows: *Luciferase-RNAi* n = 115, *Hrs-RNAi* n = 98.

### Fecundity analysis and F1 generation

Single 10-day old F0 males were crossed with 3-to 5-day old *yw* wildtype females. F0 males were removed after 20 hours and the females were placed on fresh food for egg laying and F1 generation. The female partners were flipped onto fresh food on day 2 and day 4 post mating, with the number of eggs laid on the food counted each time. Generated F1 flies were selected to carry identical balancers inherited from their fathers, so they would not carry any of the paternal *dve-Gal4* or *UAS-RNAi* components. No outliers were identified using ROUT (Q = 1 %).

The sample sizes for the fecundity analysis were as follows: number of crosses of female *yw* flies with male F0 flies expressing *Luciferase-RNAi* n = 38 and *Hrs-RNAi* n = 30 under *dve-Gal4* control.

### Bodyweight measurement of male F0 flies and their F1 offspring

Bodyweight was measured as previously described by Tennessen *et al*. [18]. F0 male flies were weighed at 10 days of age, whereas male and female F1 offspring were weighed when they were 5-days old. Briefly, 5-10 flies were weighed on an ultra-sensitive scale with up to 0.01 mg precision. To determine the average fly weight, the weight of the tube containing flies was subtracted from the weight of the tube without flies and then divided by the number of flies in the tube.

The following number of samples were removed as outliers using ROUT (Q = 1 %): male F1 from *Hrs-RNAi* fathers n = 1.

The sample sizes for the bodyweight analysis were as follows: male F0 expressing *Luciferase-RNAi* n = 36, male F0 expressing *Hrs-RNAi* n = 33; female F1 from *Luciferase-RNAi* fathers n = 18, female F1 from *Hrs-RNAi* fathers n = 25, male F1 from *Luciferase-RNAi* fathers n = 19, male F1 from *Hrs-RNAi* fathers n = 23. Each sample refers to a pool of 5-10 flies.

### Triglyceride (TAG Content) measurement of male F0 flies and their F1 offspring

To determine the TAG content, the procedure was followed as previously described by Tennessen *et al*. [18]. Male F0 flies were measured when they were 10 days old, whereas male and female F1 offspring were analysed at 5 days of age. In short, 5 flies were homogenised in 100 µl cold PBS + 0.05 % Tween 20 (PBST) and then heated for 10 min at 70°C. Following this, 20 µl PBST or 20 µl triglyceride reagent (Sigma #T2449) was added to the 20 µl fly sample. The mixtures were incubated for 30-60 min at 37°C, followed by centrifugation at full speed. The samples were transferred into a 96-well plate (30 µl) and treated with 100 µl free glycerol reagent (Sigma #F6428). After 5 min incubation at 37°C, the absorbance of the sample mixtures was measured at 540 nm on an Infinite M Plex plate reader (Tecan). The absorbance of free glycerol in PBST treated samples was subtracted from the absorbance of samples treated with triglyceride reagent. The TAG concentration for each sample was calculated using the standard curve from glycerol standards (Sigma #G7793). A Bradford assay was carried out as described by manufacturer’s instructions to measure the protein amount for normalisation.

The following sample numbers were identified as outliers and removed using ROUT (Q = 1 %): female F1 from *Hrs-RNAi* fathers n = 3.

The sample sizes for the TAG content analysis were as follows: male F0 expressing *Luciferase-RNAi* n = 35, male F0 expressing *Hrs-RNAi* n = 34; female F1 from *Luciferase-RNAi* fathers n = 28, female F1 from *Hrs-RNAi* fathers n = 25, male F1 from *Luciferase-RNAi* fathers n = 28, male F1 from *Hrs-RNAi* fathers n = 28. Each sample refers to a pool of 5 flies.

### Learning assay

Groups of 5-to 7-day-old flies were starved for 24 hours on wet KIMTECH wipes (Kimberly-Clark Worldwide Inc., UK) at 25°C and 60% humidity. For each line and gender, five replicates were included, each of which consisted of 20 – 30 flies. The test was performed using a standard T-Maze with two odours, 3-octanol (OCT) and 4-methylcyclohexanol (MCH). The odour preference test was performed separately for each line and gender and the concentration of odour was adjusted to a point where flies demonstrated no preference for any of the odours. Flies were subjected to the first odour (CS-) for two minutes. After one minute of clean air, a second odour was presented for two minutes along with dried filter paper that had been soaked in 2M sucrose (CS+). After 1 min of clean air, memory was subsequently assessed by testing flies for their odour preference between the CS- and the CS+ odours in a T-maze for 1 min. A preference index (PI) was calculated as the number of flies in the CS+ arm of the T-maze divided by the total number of tested flies: PI= CS+ / (CS+ + CS-).

### Climbing assay

For each line and gender, six replicates were included, each of which consisted of 10 flies at the age of 5-7 days. The flies of each group were placed in an empty vial with a line 5 cm from the bottom. Flies were gently tapped to the bottom of the vial and the number able to cross the line within 10 seconds was recorded. This was repeated three times for each vial with a one-minute interval, and the percentage of live flies still climbing above the line was averaged for a given line.

### RNA isolation and mRNA sequencing of F1 offspring

The gene expression profile of 5-day old male and female F1 offspring from fathers expressing *Luciferase-RNAi* and *Hrs-RNAi* under *dve-Gal4* control was analysed by mRNA sequencing. To isolate RNA, 5 flies were homogenized in TRI Reagent (Sigma Aldrich #T9424) and incubated at room temperature for 5 min. To separate the aqueous phase from the organic phase, chloroform (Carl Roth #3313.2) was added to each sample and centrifuged. Nucleotides were precipitated with isopropanol (Carl Roth #6752.2) and washed with 75% ethanol (Carl Roth #9065.2). DNA was removed by incubating the RNA samples with DNase and the Monarch RNA Cleanup Kit (New England Biolabs #T2040L) was used following the manufacturer’s instructions to improve RNA purity and quality.

For RNA sequencing, mRNA was isolated from RNA samples with NEBNext Poly(A) mRNA Magnetic Isolation Module (NEB #E7490) following the manufacturer’s instructions. LM-Seq library was prepared as previously published in [19]. In brief, RNA samples were fragmented at 80°C for 7 min and cooled on ice. For cDNA synthesis, samples were incubated with SmartScribe Reverse Transcriptase (100 U) and random hexamer oligo primer for 10 min at 23°C, then 30 min at 42°C, followed by 10 min at room temperature. Samples were treated with RNase A and RNase H for 15 min at 37°C and 5 min at 95°C to remove RNA. Purification of cDNA was carried out using Ampure Clean Up beads (Agencourt #A63881). Samples were ligated with a 5’ Illumina adaptor with an overnight incubation at 22°C with T4 RNA ligase I. Amplification and labelling of samples were carried out by PCR using FailSafe PCR enzyme and oligos that contained Illumina adaptors and index primers with unique nucleotides for each sample.

Size selection was performed using SPRIselect beads (Beckman Coulter GmbH # B23318). The cDNA concentration was quantified by a Qubit dsDNA High Sensitivity Assay Kit (Thermo Fisher Scientific # Q32851). For the library generation, 10 ng cDNA was pooled from each sample. The library quality and fragment size were quantified with an Agilent High Sensitivity DNA chip (Agilent #5067-4626) on a Bioanalyzer (2100 Bioanalyzer). Sequencing was carried out on an Illumina NovaSeq 6000 system. Fastq files were generated using bcl2fastq2 (v2.20) and the adaptor sequences were removed from the sequencing reads using CutAdapt (https://usegalaxy.org/). Trimmed reads were mapped to the Drosophila melanogaster transcriptome (dm3) using HISAT2 (v2.1.0) in Galaxy (https://usegalaxy.org/) with forward strand information and default settings.

Bamfiles were indexed with Samtools and count matrices were generated using Genomic Alignments in R. The DESeq2 package was used for the library normalisation and quantification of differentially expressed genes (DEG) between samples [20]. The significance threshold was set at an adjusted p-value < 0.05.

Sample sizes for the mRNA sequencing of F1 flies were as follows: female F1 n = 5; male F1 n = 5. Each sample refers to a pool of 5 flies.

### Gene ontology analysis

Gene Ontology (GO) analysis (GO Consortium, http://geneontology.org/) powered by PANTHER was carried out to determine enriched biological processes based on the significantly differentially expressed genes (DEG). The significance threshold was set at a false discovery rate < 0.05.

### Statistical analysis

Egg count, bodyweight (F0 and F1 samples) and TAG content (F0 and F1 samples) measurements were repeated at least three times from distinct sample batches. For each experimental batch the data was standardised to the z-score. Data was analysed in Microsoft Excel (2010) and GraphPad Prism (Version 9.3.1). Outliers were removed using ROUT (Q = 1 %). Unless otherwise specified, F0 data was analysed with an unpaired t-test with Welch’s correction or Mann-Witney t-test depending on the normal distribution assessment using the Shapiro-Wilk test. Two-way ANOVA with Tukey’s multiple comparisons test was carried out to assess statistical significance in the F1 datasets.

